# ZCCHC17 modulates neuronal RNA splicing and supports cognitive resilience in Alzheimer’s disease

**DOI:** 10.1101/2023.03.21.533654

**Authors:** Anne Marie W. Bartosch, Elliot H. H. Youth, Shania Hansen, Maria E. Kaufman, Harrison Xiao, So Yeon Koo, Archana Ashok, Sharanya Sivakumar, Rajesh K. Soni, Logan C. Dumitrescu, Tiffany G. Lam, Ali S. Ropri, Annie J. Lee, Hans-Ulrich Klein, Badri N. Vardarajan, David A. Bennett, Tracy L. Young-Pearse, Philip L. De Jager, Timothy J. Hohman, Andrew A. Sproul, Andrew F. Teich

## Abstract

ZCCHC17 is a putative master regulator of synaptic gene dysfunction in Alzheimer’s Disease (AD), and ZCCHC17 protein declines early in AD brain tissue, before significant gliosis or neuronal loss. Here, we investigate the function of ZCCHC17 and its role in AD pathogenesis. Co-immunoprecipitation of ZCCHC17 followed by mass spectrometry analysis in human iPSC-derived neurons reveals that ZCCHC17’s binding partners are enriched for RNA splicing proteins. ZCCHC17 knockdown results in widespread RNA splicing changes that significantly overlap with splicing changes found in AD brain tissue, with synaptic genes commonly affected. ZCCHC17 expression correlates with cognitive resilience in AD patients, and we uncover an APOE4 dependent negative correlation of ZCCHC17 expression with tangle burden. Furthermore, a majority of ZCCHC17 interactors also co-IP with known tau interactors, and we find significant overlap between alternatively spliced genes in ZCCHC17 knockdown and tau overexpression neurons. These results demonstrate ZCCHC17’s role in neuronal RNA processing and its interaction with pathology and cognitive resilience in AD, and suggest that maintenance of ZCCHC17 function may be a therapeutic strategy for preserving cognitive function in the setting of AD pathology.

**Significance:** Abnormal RNA processing is an important component of AD pathophysiology. We show here that ZCCHC17, a previously identified putative master regulator of synaptic dysfunction in AD, plays a role in neuronal RNA processing, and illustrate that ZCCHC17 dysfunction is sufficient to explain some of the splicing abnormalities seen in AD brain tissue, including synaptic gene splicing abnormalities. Using data from human patients, we demonstrate that ZCCHC17 mRNA levels correlate with cognitive resilience in the setting of AD pathology. These results suggest that maintenance of ZCCHC17 function may be a therapeutic strategy for supporting cognitive function in AD patients, and motivate future work examining a possible role of abnormal RNA processing in AD-related cognitive decline.

## Introduction

Synaptic dysfunction linked to cognitive performance is an early occurrence in AD animal models (Trinchese et al., 2004; Smith et al., 2009; Siskova et al., 2014), and dysregulation of synaptic gene expression has consistently been shown in AD autopsy tissue (Loring et al., 2001; Colangelo et al., 2002; Liang et al., 2008). Understanding the molecular basis of synaptic dysfunction is therefore of high importance in the AD field (Teich et al., 2015). We previously used mutual information relationships between RNA expression profiles of autopsy brain tissue to identify transcriptional regulators that may be driving dysregulated synaptic gene transcription in AD, and reported ZCCHC17 as a candidate regulator that is predicted to have reduced activity in AD, leading to dysregulation of gene expression across a broad range of categories (including synaptic) (Tomljanovic et al., 2018). ZCCHC17 was discovered in 2002 while screening a cDNA library for RNA binding proteins (Gueydan et al., 2002) and was also independently found in 2003 using a yeast two-hybrid screen that searched for pinin-interacting proteins (Chang et al., 2003). Additionally, ZCCHC17 interacts with SRrp37, which regulates splicing of pre-mRNA (Ouyang, 2009), and has recently been shown to interact with the splicing factors SRSF1 and SRSF2 (Lin et al., 2017). ZCCHC17 contains an S1 RNA binding domain, zinc-finger (CCHC) domain, and 2 nuclear localization signals, and northern blot analysis has shown that its transcripts are found throughout the body, with strongest expression in the normal brain, heart, skeletal muscle, and thymus (Chang et al., 2003). Although the exact function of ZCCHC17 is unknown, recent evidence showing that it is involved in mRNA (Lin et al., 2017) and rRNA (Lin et al., 2019) processing have led some to suggest that it may help coordinate a wide range of homeostatic functions in the cell (Lin et al., 2017). Our prior work demonstrated that ZCCHC17 protein is expressed in neurons and declines early in AD brain tissue before significant gliosis or neuronal loss, while knockdown of ZCCHC17 in rat neuronal cultures leads to dysregulation of a wide range of genes, including synaptic genes (Tomljanovic et al., 2018). ZCCHC17 has been bioinformatically linked to AD more generally by others as well (Li et al., 2015). However, the activity of ZCCHC17 in human neurons has not yet been explored, and its clinical relevance in AD patients has not been examined.

Here, we investigate the functional role of ZCCHC17 in a human iPSC-derived neuronal model, and explore how this role may contribute to clinical decline in AD patients. By performing co-immunoprecipitation of ZCCHC17 in human neurons followed by mass spectrometry analysis, we show that ZCCHC17 interacts with a network of splicing proteins. Furthermore, ZCCHC17 knockdown leads to dysregulation of mRNA splicing across a broad range of gene categories (including synaptic), and these differentially spliced genes overlap significantly with those seen in AD brain tissue. ZCCHC17 expression correlates with cognitive resilience in AD patients through two different approaches (using ROSMAP data and PrediXcan analysis), and we uncover an APOE4-dependent correlation of ZCCHC17 expression with neurofibrillary tangle burden. Finally, we show that a majority of ZCCHC17 interactors identified by co-IP also co-IP with known tau interactors, and we find significant overlap between alternatively spliced genes in ZCCHC17 knockdown and tau overexpression neurons. Taken together, the above results support a role for ZCCHC17 in neuronal RNA splicing and cognitive resilience in AD, and point to tau as a possible mediator of ZCCHC17 dysfunction.

## Methods

### hiPSC and Neuronal Cell Culture

All human cell line work was performed on de-identified cell lines and approved by the Columbia University Institutional Review Board (IRB). IMR90 cl.4 hiPSCs (WiCell) (Yu et al., 2007; Yu et al., 2009; Chen et al., 2011; Hu et al., 2011) were grown in StemFlex media (Thermo Scientific) on Cultrex (Biotechne) and passaged with ReLeSR as described previously (Song et al., 2021). Human neural progenitor cells (NPCs) were generated using dual SMAD inhibition as non-adherent embryoid bodies, followed by plating on polyornithine (10 ug/mL, Sigma P4957)/laminin (10 ug/mL, R&D Systems 3400-010-02) and subsequent manual rosette selection and expansion, as described previously (Topol et al., 2015; Sun et al., 2019). NPCs were maintained on Matrigel (Corning 354230) and split 1:2 to 1:3 at every 5-7 days. Neuronal differentiations were carried out by plating 200,000 cells/12 well-well or 500,000 cells/6 well-well in DMEM/F12 base media (Gibco 11320-033) supplemented with B27 (Gibco 17504-044), N2 (Gibco 17502-048), penicillin/streptomycin (Gibco 15140-122), BDNF (20 ng/mL, R&D Systems 248-BDB), and laminin (1 ug/mL). After 1 week of differentiation, AraC (Tocris 4520) was added at 100 nM to reduce remaining NPC proliferation. Neurons were differentiated for 6-11 weeks post NPC stage.

### Lentiviral Construct and Lentiviral Particle Formation

cDNA corresponding to human ZCCHC17 transcript variant 3 was ordered from the MGC Collection from Dharmacon (Horizon Discovery Biosciences MHS6278-202760302). This was used as a template for PCR using Amplitag Gold (Life Technologies, as per user instructions), utilizing the following primers that introduced a Kozak sequence and an N-terminal FLAG sequence after the translational start site: <u>ZCC_NFLAG_TOPO_F</u>:CACCATGGACTACAAAGACGATGACGACAAGAATTCAGGAAGGCCTGAGACC

<u>ZCC_NFLAG_TOPO_R:TCACTCCTTGTGCTTCTTCTTGTGC</u>.

Purified PCR products were then TOPO-cloned into pENTRE/D-TOPO (Life Technologies). Transformed bacterial colonies (Stb3, Life Technologies) were miniprepped (Qiagen) and confirmed to be correct by Sanger sequence (Genewiz/Azenta). One correct clone was subsequently Gateway-cloned (Life Technologies, as per user instructions) into pLEX-305, which was a gift from David Root (Addgene plasmid#41390; http://n2t.net/addgene:41390; RRID:Addgene_41390).

Transformed bacterial colonies were miniprepped and confirmed to be correct by Sanger sequence, and then midiprepped (Machery-Nagel) for use in lentiviral particle generation. A similar strategy was later used for C-terminal tagged ZCCHC17, using the following primers:

<u>ZCC_CFLAG_TOPO_F</u>:CACCATGAATTCAGGAAGGCCTGAGACC <u>ZCC_CFLAG_TOPO_R</u>:TCACTTGTCGTCATCGTCTTTGTAGTCCTCCTTGTGCTTCTTCTTGTGC.

“True empty” pLEX-305 was generated by excision of the ccdB8 selection cassette using Age1 and Xho1 restriction digest, blunting via Large Klenow Fragment, and gel purification of the digested plasmid (2243 bp excised), followed by blunt ligation and bacterial transformation. After bacterial colonies were miniprepped and confirmed to be correct by Sanger sequence, a correct clone was midiprepped for lentiviral particle production.

Lentiviral particles from N- and C-terminal FLAG-tagged ZCCHC17 as well as empty pLEX_305 vector were generated as described previously (Song et al., 2021). Briefly, HEK 293T cells were plated at 180,000 cells/cm^2^ on gelatin-coated 15-cm plates and fed with 20 ml FM10 media. Cells were confirmed to be 80-90% confluent preceding transfection the following day. 80 µl Lipofectamine 3000 (Invitrogen L3000-015) was mixed with 2 ml Opti-MEM (Gibco 31985-070) in a 15-ml conical tube and vortexed for 3 seconds. A plasmid DNA master mix was prepared in a separate 15-ml conical tube, comprising 20 µg FLAG-ZCCHC17, ZCCHC17-FLAG, or control pLEX_305 empty vector, 15 µg psPAX2 (Addgene #12260), and 10 µg VSV-G (Addgene #14888) in 2 ml Opti-MEM, supplemented with 80 µl P3000 reagent (Invitrogen L3000-015). The 2 mixes were combined and incubated at room temperature for 10 minutes, and the transfection mixture was subsequently pipetted onto cells in a dropwise manner. Cells were then incubated for 5-6 hours at 37°C, after which the transfection media was replaced with 18.5 ml fresh FM10. Viral supernatants were collected 48 and 72 hours post-transfection and centrifuged for 3 min at 300 × g. Supernatants were then combined, vacuum-filtered, transferred to new tubes, supplemented with 1/3 volume Lenti-X Concentrator (Takara 631232), mixed by gentle inversion, and incubated overnight at 4°C. The following day, samples were centrifuged for 45 minutes at 4°C at 1,500 × g. Supernatants were removed and the lentiviral pellets were resuspended in DPBS (Gibco 14190-044) and stored at -80°C.

### Immunofluorescence

hiPSC-derived neurons on glass coverslips in 12-well plates were infected with FLAG-tagged ZCCHC17 and control lentivirus one week prior to fixation and immunostained to assess expression of constructs via immunofluorescence. Cells were fixed in 4% paraformaldehyde for 20 minutes, incubated with 0.2% Triton X-100 (Thermo 85111) in PBS for 5 minutes, and washed 3x with PBS. Coverslips were incubated for 1 hour at room temperature in blocking solution, consisting of 5% goat serum in PBS, and then overnight at 4°C in primary antibody solution, consisting of anti-ZCCHC17 (1:1000, Abcam ab80454) and anti-FLAG (1:1000, Sigma F3165) in blocking solution. Coverslips were rinsed 3x with PBS, incubated with 1:1000 secondary antibody in blocking solution for 1 hour at room temperature, and washed 3x with PBS. Coverslips were incubated for 5 minutes with DAPI in PBS, washed 1x with PBS, mounted with Vectashield (Vector H-1000), and stored at 4°C. Imaging was carried out using an LSM 800 confocal microscope (Zeiss).

### Immunoprecipitation

hiPSC-derived neurons were differentiated in 6-well plates and infected with FLAG-tagged ZCCHC17 and control lentivirus one week prior to immunoprecipitation. Cells were rinsed 2x with cold PBS, incubated on ice for 15 minutes with cold IP lysis buffer (Thermo 87787) supplemented with protease and phosphatase inhibitors (Thermo 1861280), collected into tubes by scraping, and centrifuged at 12,000 × g for 10 minutes. 40 uL/sample of anti-FLAG conjugated magnetic bead slurry (Sigma M8823) was prepared by washing 6x with PBS to remove glycerol from beads. Lysates were diluted to 1 mL total volume with additional IP lysis buffer with inhibitors. Anti-FLAG beads were added to samples, gently mixed, and incubated at room temperature on a rotator for 2 hours. Anti-FLAG beads with bound sample were magnetically separated from remaining unbound sample and rinsed 2x with 1 mL PBS. Immunoprecipitated samples were further processed by *in-gel* digestion for mass spectrometry or were prepared for western blot as detailed below.

### Western Blotting

Tris-glycine SDS sample buffer (Thermo LC2676) was added to beads with bound sample and boiled for 3 min. Bound sample in sample buffer was magnetically separated from beads and stored at -80°C. Thawed samples were supplemented with sample reducing agent (Thermo NP0009), and boiled for 5 min prior to loading to 4-20% gradient gels (Thermo XP04200BO). Samples were separated by SDS-PAGE and transferred to nitrocellulose membrane at 30V overnight at 4°C. Membranes were incubated for 1 hour in 100% sea block blocking buffer (Thermo 37527) on a rocker at room temperature. Anti-FLAG (1:1000, Sigma F3165), anti-ZCCHC17 (1:1000, Abcam ab80454), anti-AP2A1 (1:1000, Proteintech 11401-1-AP) or anti-hnRNPU (1:1000, Abclonal A3917) were added to 50% sea block in TBS-Tween (TBST) buffer solution and incubated at 4°C overnight with rocking. After washing with TBST, secondary antibodies (Licor) were added at 1:10,000 to 50% sea block in TBST and incubated for 1 hour at room temperature with rocking. Membranes were washed and imaged using the LI-COR Odyssey imaging system.

### *In-gel* Digestion for Mass Spectrometry

Immunoprecipitated samples (n = 3 per group, including FLAG-ZCCHC17, ZCCHC17-FLAG, and negative control samples) were separated on 4-12% gradient SDS-PAGE and stained with SimplyBlue. Protein gel slices were excised and *in-gel* digestion was performed as previously described (Shevchenko et al., 2006), with minor modifications. Each gel slice was washed with 1:1 Acetonitrile and 100 mM ammonium bicarbonate for 30 minutes, then dehydrated with 100% acetonitrile for 10 minutes until shrunk. The excess acetonitrile was then removed and the slice was dried in a speed-vacuum at room temperature for 10 minutes. The gel slice was reduced with 5 mM DTT for 30 min at 56 °C in an air thermostat, cooled down to room temperature, and alkylated with 11 mM IAA for 30 minutes with no light. The gel slice was then washed with 100 mM of ammonium bicarbonate and 100% acetonitrile for 10 minutes each. Excess acetonitrile was removed and dried in a speed-vacuum for 10 minutes at room temperature, after which the gel slice was re-hydrated in a solution of 25 ng/μl trypsin in 50 mM ammonium bicarbonate for 30 minutes on ice and digested overnight at 37 ^0^C in an air thermostat. Digested peptides were collected and further extracted from the gel slice in extraction buffer (1:2 ratio by volume of 5% formic acid:acetonitrile) by shaking at high speed in an air thermostat. The supernatants from both extractions were then combined and dried in a speed-vacuum. Peptides were dissolved in 3% acetonitrile/0.1% formic acid.

### Liquid Chromatography with Tandem Mass Spectrometry (LC-MS/MS)

Desalted peptides were injected in an EASY-Spray^™^ PepMap^™^ RSLC C18 50cm X 75cm ID column (Thermo Scientific) connected to an Orbitrap Fusion^™^ Tribrid^™^ (Thermo Scientific). Peptide elution and separation was achieved at a non-linear flow rate of 250 nl/min using a gradient of 5%-30% of buffer B (0.1% (v/v) formic acid, 100% acetonitrile) for 110 minutes, with column temperature maintained at 50 °C throughout the entire experiment. The Thermo Scientific Orbitrap Fusion Tribrid mass spectrometer was used for peptide tandem mass spectroscopy (MS/MS). Survey scans of peptide precursors were performed from 400 to 1500 *m/z* at 120K full width at half maximum (FWHM) resolution (at 200 *m/z*) with a 2 × 10^5^ ion count target and a maximum injection time of 50 ms. The instrument was set to run in top speed mode with 3-second cycles for the survey and the MS/MS scans. After a survey scan, MS/MS was performed on the most abundant precursors, i.e., those exhibiting a charge state from 2 to 6 of greater than 5 × 10^3^ intensity, by isolating them in the quadrupole at 1.6 Th. We used collision-induced dissociation (CID) with 35% collision energy and detected the resulting fragments with the rapid scan rate in the ion trap. The automatic gain control (AGC) target for MS/MS was set to 1 × 10^4^ and the maximum injection time was limited to 35ms. The dynamic exclusion was set to 45s with a 10 ppm mass tolerance around the precursor and its isotopes. Monoisotopic precursor selection was enabled.

### LC-MS/MS Data Analysis

Raw mass spectrometric data were analyzed using the MaxQuant environment (v1.6.1.0) and Andromeda for database searches (Cox et al., 2011) at default settings with a few modifications. The default was used for first search tolerance and main search tolerance (20 ppm and 6 ppm, respectively). MaxQuant was set up to search with the reference human proteome database downloaded from Uniprot (https://www.uniprot.org/proteomes/UP000005640). MaxQuant performed the search for trypsin digestion with up to 2 missed cleavages. Peptide, site and protein false discovery rates (FDR) were all set to 1% with a minimum of 1 peptide needed for identification; label-free quantitation (LFQ) was performed with a minimum ratio count of 1. The following modifications were used for protein quantification: oxidation of methionine (M), acetylation of the protein N-terminus, and deamination for asparagine or glutamine (NQ). Results obtained from MaxQuant were further analyzed using R. Protein identifications were filtered for common contaminants. Proteins were considered for quantification only if they were found in at least two replicate samples from a test group. Significant enrichment in protein abundance was determined by t-test with a significance threshold of adjusted p-value < 0.1 (permutation-based FDR correction) and log_2_(FoldChange) > 0.3.

### ZCCHC17 siRNA Delivery

hiPSC-derived neurons were fed and treated with 1 μM Accell self-delivering siRNA, consisting of a pool of four unique ZCCHC17 sequences (Horizon Discovery Biosciences E-105851-00) or four non-targeting control sequences (Horizon Discovery Biosciences D-001910-10). Cells were harvested 7 days after siRNA treatment for corresponding experiments.

### RNA Extraction

hiPSC-derived neurons were lysed in Qiazol lysis reagent (Qiagen 79306) on ice and frozen at -80°C. 12 samples were collected for RNA sequencing (n = 6 ZCCHC17 knockdown biological replicates and n = 6 negative control biological replicates). RNA was extracted using the miRNeasy Mini kit (Qiagen 217004). RNA library preparation and sequencing were conducted at GENEWIZ, LLC (South Plainfield, NJ, USA) as described below.

### Library Preparation with Stranded PolyA Selection

Total RNA samples were quantified using a Qubit 2.0 Fluorometer (Life Technologies, Carlsbad, CA, USA) and RNA integrity was checked by an Agilent TapeStation 4200 (Agilent Technologies, Palo Alto, CA, USA). RNA sequencing libraries were prepared using the NEBNext Ultra II Directional RNA Library Prep Kit for Illumina, following the manufacturer’s instructions (NEB, Ipswich, MA, USA). Briefly, mRNAs were first enriched with Oligo(dT) beads, and enriched mRNAs were fragmented for 15 minutes at 94 °C. First strand and second strand cDNAs were subsequently synthesized. cDNA fragments were end repaired and adenylated at their 3’ ends, and universal adapters were ligated to the cDNA fragments, followed by index addition and library enrichment by limited-cycle PCR. The sequencing libraries were validated on an Agilent TapeStation (Agilent Technologies, Palo Alto, CA, USA), and quantified using a Qubit 2.0 Fluorometer (Invitrogen, Carlsbad, CA) as well as by quantitative PCR (KAPA Biosystems, Wilmington, MA, USA).

### HiSeq Sequencing

The sequencing libraries were pooled and clustered on 4 lanes of a flowcell. After clustering, the flowcell was loaded on an Illumina HiSeq instrument (4000 or equivalent) according to the manufacturer’s instructions. The samples were sequenced using a 2×150bp paired-end (PE) configuration. Image analysis and base calling was performed by the HiSeq Control Software (HCS). Raw sequence data (.bcl files) generated from Illumina HiSeq was converted into fastq files and de-multiplexed using Illumina’s bcl2fastq 2.17 software. One mismatch was allowed for index sequence identification.

### RNA Sequencing Data Preprocessing

Raw RNA-seq data for ZCCHC17 knockdown neurons, Religious Orders Study and Rush Memory and Aging Project (ROSMAP) bulk dorsolateral prefrontal cortex (DLPFC) tissue (Bennett et al., 2018; Mostafavi et al., 2018), and tau overexpression neurons (Raj et al., 2018) was preprocessed as follows. The quality of all FASTQ files was assessed using FastQC (v0.11.9), and all samples for which one or both paired-end FASTQ files failed the “per base sequencing quality” metric were excluded from downstream analysis in order to ensure confidence in sequence-based inferences. Reads were aligned to the GRCh38 genome (Ensembl Release 101) using STAR (v2.7.6a) in 2-pass mapping mode with standard ENCODE parameters. Gene counts were quantified from STAR-aligned BAM files using featureCounts (v2.0.1). Only samples with RIN ≥ 5 were used for downstream analysis.

### RNA Splicing Analysis

Differential splicing analysis was performed using LeafCutter (v0.2.9) in Python (v2.7) and R (v3.4.4) (Li et al., 2018). LeafCutter enables annotation-agnostic quantification of RNA splicing by grouping introns inferred from aligned reads into splicing clusters which enable statistical comparison of differential usage between 2 groups. In brief, splicing events were extracted from STAR-aligned BAM files using regtools (v0.5.2) with minimum anchor length 8 bp, minimum intron length 20 bp, and maximum intron length 1M bp. The resulting junction files were then clustered with LeafCutter, allowing introns of maximum length 1M bp and requiring each intron to be supported by at least 100 reads and account for at least 5% of the reads in its splicing cluster. Differential splicing analysis was performed by LeafCutter, requiring that each intron analyzed be supported by at least 30 reads in all samples (for NPC-derived neurons and tau overexpression neurons) or in at least half of the samples in each clinical diagnosis group (for ROSMAP bulk DLPFC tissue). Differential splicing in ROSMAP was assessed between AD samples (defined as those with a clinical diagnosis of Alzheimer’s dementia and no other cause of cognitive impairment) and control samples (defined as those with no cognitive impairment on clinical assessment), controlling for covariates including study (ROS or MAP), sequencing batch and depth, donor age and sex, PMI, and RIN. Downstream analysis of differential splicing results was performed in R (v4.0.2). Differentially spliced clusters were identified by FDR-corrected p-value < 0.05. Differentially spliced genes (DSGs) were inferred from differentially-spliced clusters which corresponded unambiguously to a single annotated gene. Individual splice junction plots were generated using LeafViz (v0.1.0). Intron usage plots were generated using ggplot2 (v3.3.2).

### Gene Ontology Analysis

DSGs were extracted from LeafCutter analysis results as described above. Gene ontology (GO) terms spanning GO levels 2-5 and all 3 GO families (biological process, molecular function, and cellular component) were characterized using the “over-representation analysis” function of the ConsensusPathDB web tool. Significantly-enriched GO terms were identified by FDR-corrected p-value < 0.05.

### Nonsense-Mediated Decay Analysis

Differential isoform analysis was performed using the IsoformSwitchAnalyzeR package (v1.10.0) in R (v4.0.2). In brief, transcript counts were quantified from STAR-aligned BAM files using Salmon (v1.4.0), and differential isoform usage was assessed using DEXSeq. Differential isoform usage in ROSMAP samples was assessed controlling for covariates including study (ROS or MAP), sequencing batch and depth, donor age and sex, PMI, and RIN. Premature termination codons (PTCs) were defined as annotated STOP codons located at least 50 bp upstream of the final canonical exon-exon junction for a given transcript, and isoforms containing PTCs were classified as NMD-sensitive. For each dataset analyzed, significant changes in differential isoform fraction (dIF) were assessed using two-sided t-tests, and associations between PTC status and dIF sign were assessed using Fisher’s exact test. NMD sensitivity plots were generated using ggplot2 (v3.3.2).

### Gene Expression Correlational Analysis

ROSMAP sample metadata including APOE genotype, cognitive metrics, and midfrontal cortex AD pathology metrics were obtained from the Rush Alzheimer’s Disease Center (Bennett et al., 2018; De Jager et al., 2018). All downstream analysis was performed in R (v4.0.2). After filtering out low-expression genes (defined as all those with <5 raw read counts in at least 90% of all samples), raw gene counts were normalized and variance-stabilized using DESeq2 (v1.28.1). The effects of sequencing batch and depth, donor age and sex, PMI, and RIN were subsequently regressed out using limma::removeBatchEffect (v3.44.3). Processed ZCCHC17 expression values were then Spearman-correlated to cognitive and AD pathology metrics using Hmisc::rcorr (v4.4.2). Separate analyses were run on the subset of samples with documented APOE genotypes to assess correlations for subgroups with and without the APOE4 risk factor allele. Correlational p-values were BH-adjusted within each analysis.

### Predicted Gene Expression Association with Resilience

We leveraged data from a published genome-wide association study (GWAS) of resilience to AD neuropathology (Dumitrescu et al., 2020), defined as better-than-predicted cognitive performance given an individual’s amyloid burden. PrediXcan (Gamazon et al., 2015) was used to quantify predicted levels of ZCCHC17 expression leveraging the GTEx database for model building and applied using GWAS data. Tissue-specific expression models were built leveraging elastic-net regression in the cis gene region (within 1Mb) and selected based on five-fold cross-validation as previously described (Gamazon et al., 2015). To determine whether genetic regulation of ZCCHC17 was related to resilience, we quantified the association between predicted ZCCHC17 expression with our published Combined Resilience trait (n = 5,108) (Dumitrescu et al., 2020) covarying for age and sex. The false discovery rate procedure was used to correct for multiple comparisons. The Generalized Berk-Jones test was used to summarize the association across tissues into a single test statistic. Secondary analyses were performed removing individuals with AD from the analytical model (n = 3,818). Finally, for all cross-validated ZCCHC17 models that related to resilience, we also assessed whether genetic regulation predicted observed transcript expression of ZCCHC17 in three brain regions, leveraging the genotype and RNA sequencing data from ROSMAP. All validation models were limited to non-Hispanic white participants without AD (n = 172) to align with the prediction models built in GTEx. For our GWAS analysis, nominally significant SNPs (*P* < 1 × 10^−5^) associated with AD were highlighted from summary statistics from (Jansen et al., 2019), which uses GRCh37.

## Results

### Identification of Binding Partners of ZCCHC17

ZCCHC17 has been studied in non-neuronal tissues and has been shown to play a diverse range of regulatory roles in RNA processing (Lin et al., 2017; Lin et al., 2019). Its function has not been previously studied in human neurons, and it may exert its regulatory effect through its CCHC-type zinc finger domain, the RNA-binding capacity of its S1 domain, or via its interactions with proteins involved in RNA processing (**Figure 1A**; note that it may be less likely to act as a classical zinc finger transcription factor as it only contains a single zinc finger domain (Lambert et al., 2018)). Since protein function is directly related to protein-protein interactions, we first performed immunoprecipitation-mass spectrometry to identify the binding partners of ZCCHC17 and elucidate its function. N- and C-terminal FLAG-tagged ZCCHC17 lentiviral constructs were expressed in human iPSC-derived neurons to enable isolation of ZCCHC17 protein and its interactors by FLAG immunoprecipitation (**Supplemental Figures S1-2**). Immunoprecipitated proteins were identified using mass spectrometry. Significant binding partners were identified by comparing proteins bound to FLAG-tagged ZCCHC17, immunoprecipitated from either FLAG-ZCCHC17- or ZCCHC17-FLAG-expressing neurons, to proteins nonspecifically bound to FLAG beads in control neurons expressing an empty vector (**Figure 1B-C**). FLAG tag location (C-terminal or N-terminal) had only a marginal effect on binding partner hits; there was a significant change in the abundance of only one protein, FTL, which binds to N-terminal FLAG-tagged ZCCHC17 to a higher degree than to C-terminal FLAG-tagged ZCCHC17. This suggests that the C-terminal FLAG tag may have generated some steric hindrance which impeded FTL’s ability to bind to ZCCHC17, although FTL remained a significant binding partner in both experiments when compared to control samples.

**Fig. 1.**
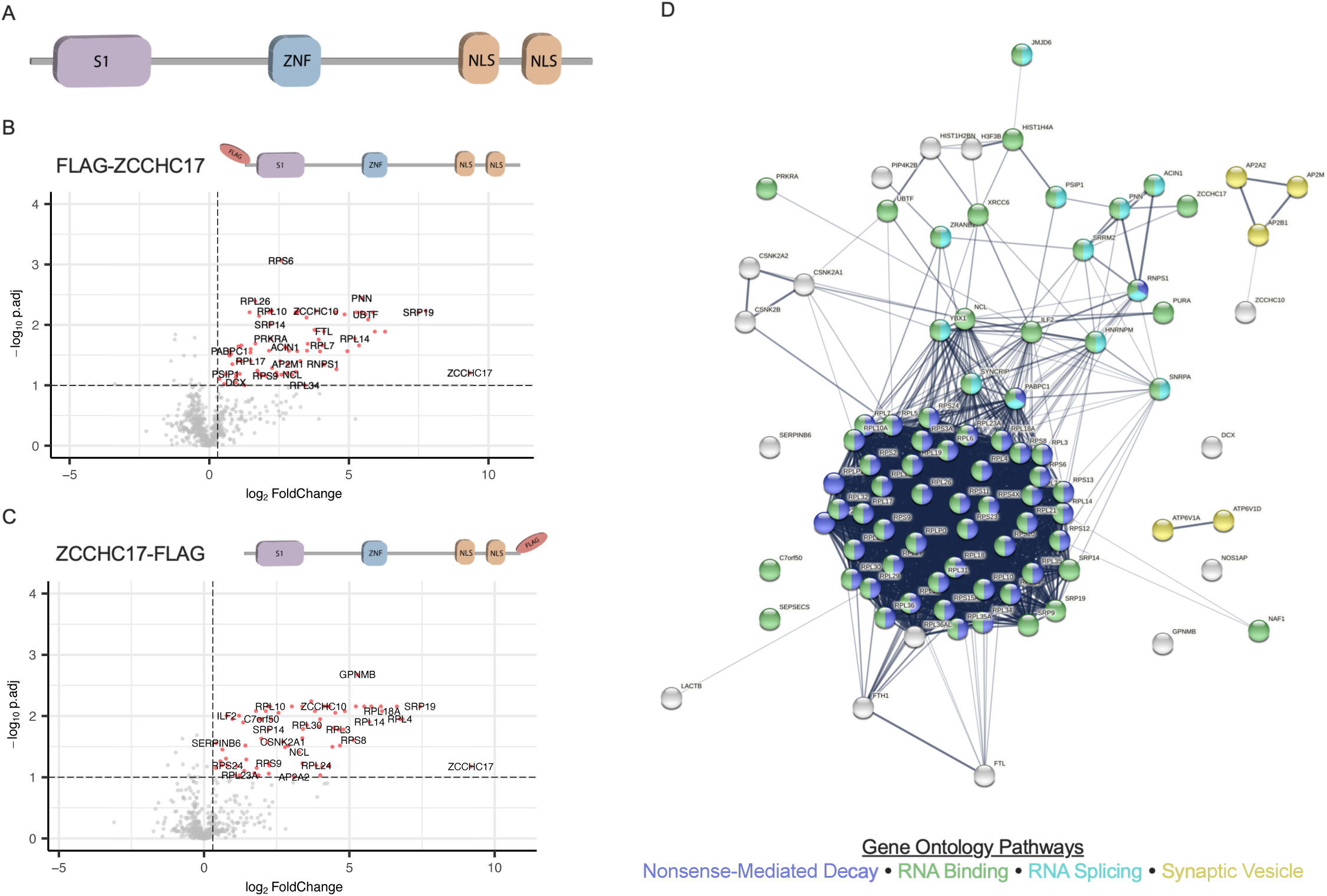
ZCCHC17 Binding Partners Include RNA Splicing, Binding, and Processing Proteins. **(A)** ZCCHC17 protein domains include an S1 RNA-binding domain, a CCHC-type zinc finger (ZNF) domain, and two nuclear localization signal (NLS) domains. (**B-C**) LC-MS/MS of bound proteins from anti-FLAG immunoprecipitation performed on human iPSC-derived neurons differentiated for 75 days post-NPC expressing N- or C- terminal FLAG-tagged ZCCHC17 compared to control neurons. Volcano plots display log_2_-fold changes and FDR-corrected p-values defined with respect to protein abundance in control neurons (n = 3 for all groups); significant hits (adjusted p < 0.1 and log_2_FC > 0.3) are highlighted in red. (**D**) STRING-derived known and predicted protein-protein interactions among ZCCHC17 binding partners, with relevant strongly-enriched gene ontology pathways highlighted. ZCCHC17 binding partners were defined as the union of significant hits from both LC-MS/MS experiments and comprise 91 unique proteins in total.

91 unique binding partners for ZCCHC17 were identified in human iPSC-derived neurons (**Figure 1D, Supplemental Table S1**). Seven of these proteins have been previously identified as ZCCHC17 binding partners in non-neuronal cells, including PNN (Chang et al., 2003; Huttlin et al., 2017; Lin et al., 2017), UBTF (Lin et al., 2019), JMJD6, FTL, SRRM2, NOS1AP, and RNPS1 (Huttlin et al., 2017), suggesting conserved interactions across cell types. Several of ZCCHC17’s binding partners have previously been highlighted for their importance in AD, including those involved in autophagy/lysosome function (AP2A1, AP2A2, ATP6V1A) (Raj et al., 2018; Wang et al., 2021a), stress granules (PABPC1, PABPC3) (Anderson and Kedersha, 2008), TIA-1 interactors (RPL6, RPL7, RPL7A, RPL10A, RPL13, RPS4X, AP2A1, AP2B1, ATP6V1A, PURA, SYNCRIP, PABPC1) (Vanderweyde et al., 2016), and RNA binding (ACIN1, SNRPB2) in the presence of AD pathology (Apicco et al., 2019). AP2A1 and HNRNPU were also shown by western blot following immunoprecipitation of FLAG-tagged ZCCHC17 (**Supplemental Figure S3**).

Enrichment analysis was performed using STRING (v11.0b), which integrates well-known classification systems including KEGG, Reactome, and Gene Ontology (**Figure 1D**) (Szklarczyk et al., 2019). Significantly-enriched gene ontology (GO) terms included numerous RNA processing pathways such as RNA binding, RNA splicing, and nonsense-mediated decay (NMD) (strength = 1.0, 0.8, 1.9; FDR-adjusted p-values for enrichment = 4.4 × 10^−51^, 5.0 × 10^−5^, 1.0 × 10^−72^, respectively). Several of ZCCHC17’s binding partners were also involved in the synaptic vesicle cycle (strength = 1.2; FDR-adjusted p-value for enrichment = 5.5 × 10^−3^).

### ZCCHC17 Knockdown Induces Differential Splicing that Overlaps with Known AD Splicing Abnormalities

The prevalence of RNA splicing proteins among ZCCHC17’s binding partners inspired further investigation of RNA splicing alterations in hiPSC-derived neurons. Thus, we treated neurons with siRNAs targeting ZCCHC17 or non-targeting control siRNAs. ZCCHC17 was the most significantly downregulated of all differentially expressed genes (DEGs), with log_2_FC = -2.5 (**Supplemental Table S2**), confirming robust knockdown (>80%). Alternative splicing analysis, conducted using the LeafCutter algorithm (Li et al., 2018), identified 732 differentially spliced intron clusters corresponding to 637 unique differentially spliced genes (DSGs) in the ZCCHC17 knockdown model (**Figure 2A, Supplemental Table S3**). Gene Ontology (GO) enrichment analysis revealed 69 significant GO terms enriched among ZCCHC17 knockdown DSGs, of which 11 were synaptic-related pathways (**Supplemental Table S3**). Top GO terms included “postsynaptic specialization organization” and “postsynaptic density organization” (adjusted p-values = 7.8 × 10^−3^ and 9.3 × 10^−3^, respectively). Taken together, this suggests that ZCCHC17 affects splicing in a significant number of genes, among which synaptic targets are enriched.

**Fig. 2.**
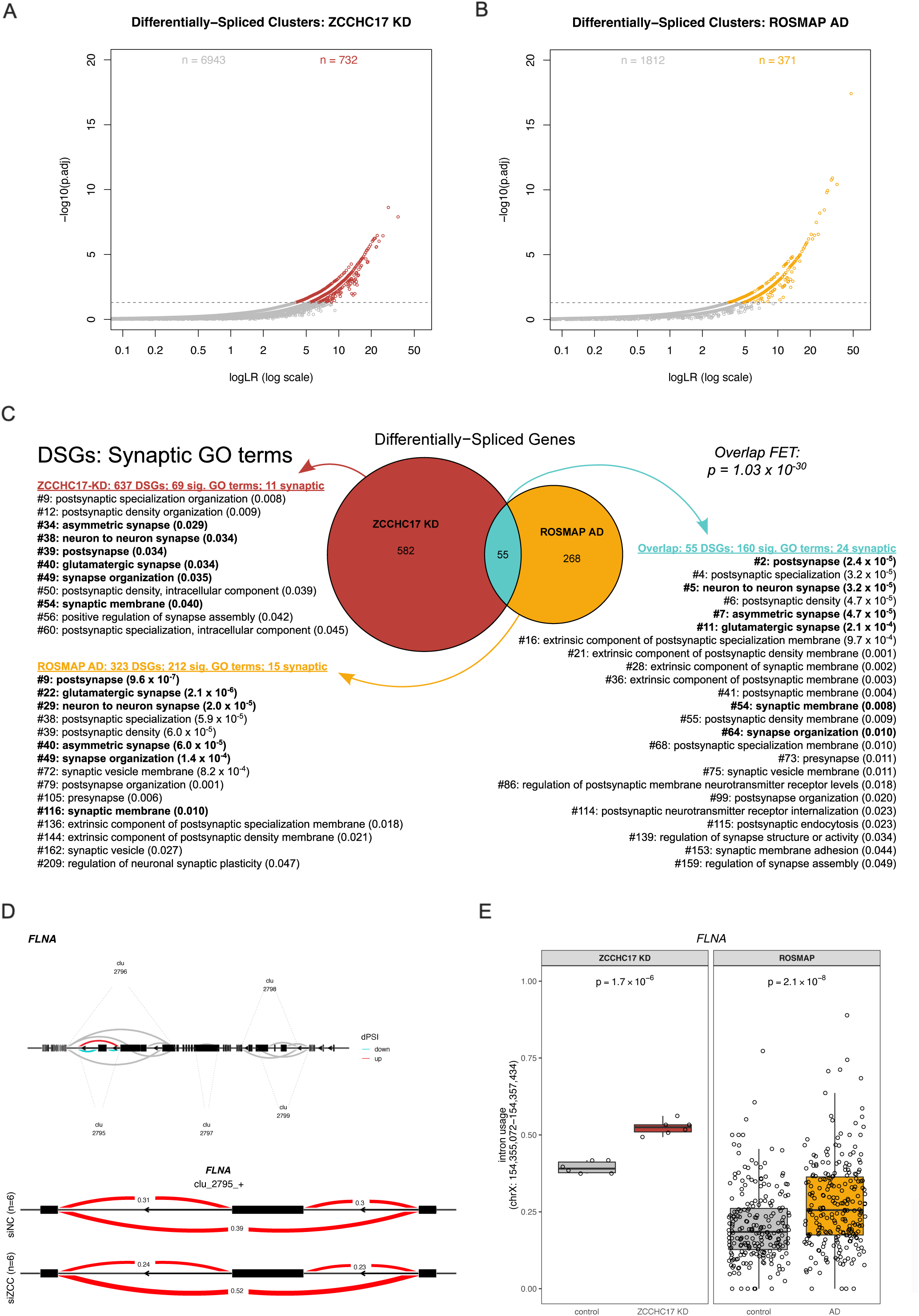
ZCCHC17 Knockdown Induces RNA Splicing Alterations in AD-Related Genes. **(A)** 732 intron clusters exhibited differential splicing between ZCCHC17 knockdown (n = 6) and controls (n = 6) in human iPSC-derived neurons differentiated for 42 days post-NPC. (**B**) 371 intron clusters exhibited differential splicing between AD and control human DLPFC tissue samples from ROSMAP. (**C**) Differentially-spliced genes (DSGs) in both ZCCHC17 knockdown neurons and AD human brain tissue are enriched for synaptic genes. There is a significant overlap in DSGs between the two datasets (p = 1.0 × 10^−30^ by Fisher’s exact test). The 55 genes that are differentially spliced in both datasets are highly enriched for synaptic genes. (**D**) In ZCCHC17 knockdown neurons, FLNA is differentially spliced at cluster 2795 (p = 2.4 × 10^−9^), corresponding to an exon skipping event. All FLNA intron clusters identified by LeafCutter are shown in the top diagram. DeltaPSI (dPSI) is the difference in usage proportion between the two groups and is red or blue for introns with significantly increasing or decreasing dPSI, respectively. Clusters shown in grey did not change significantly. In the lower diagram, exons in cluster 2795 are shown in black, with excision proportions for introns in both experimental groups shown in red. (**E)** Exon skipping occurs at the same location and to a similar degree in both ZCCHC17 knockdown neurons and AD human brain tissue, with significant changes in both contexts (p = 1.7 × 10^−6^ and 2.1 × 10^−8^, respectively).

We have previously demonstrated that ZCCHC17 levels are reduced early in AD, before significant astrogliosis or neuronal loss (Tomljanovic et al., 2018). To identify whether ZCCHC17 might alter the RNA processing of any synaptic genes affected in AD, differential splicing analysis was repeated using ROSMAP data comprising postmortem DLPFC brain tissue samples from 238 AD and 227 control patients (Raj et al., 2018). 371 differentially spliced intron clusters were identified in ROSMAP AD samples, corresponding to 323 unique differentially spliced genes (**Figure 2B, Supplemental Table S3**). GO enrichment analysis revealed 212 significant GO terms enriched among ROSMAP AD DSGs, of which 15 were synaptic-related pathways (**Supplemental Table S3**). “Postsynapse” was a top GO term (adjusted p-value = 9.6 × 10^−7^).

When ZCCHC17 knockdown and ROSMAP AD splicing analyses were compared, 55 DSGs emerged in common (**Figure 2C**). This represents 17% of all DSGs we detected in the bulk DLPFC RNA-seq data from ROSMAP, and is statistically significant (p = 1.0 × 10^−30^by Fisher’s exact test). GO enrichment analysis of the 55 shared DSGs revealed 160 significant GO terms, of which 24 were synaptic-related pathways (**Supplemental Table S3**). Top GO terms enriched among overlapping DSGs included “postsynapse,” “postsynaptic specialization,” “neuron to neuron synapse,” and “postsynaptic density” (adjusted p-values = 2.4 × 10^−5^, 3.2 × 10^−5^, 3.2 × 10^−5^, and 4.7 × 10^−5^, respectively). Examination of individual shared DSGs revealed similar patterns of differential splicing induced by ZCCHC17 knockdown and presence of AD. *FLNA* was the most dramatically affected synaptic DSG in both contexts; it was the most significant hit in the ZCCHC17 knockdown neuronal model and the third-most significant hit in the ROSMAP AD analysis (**Supplemental Table S3, Figure 2D**). Filamin A (FLNA) is the most common member of the filamin family of proteins, which bind to actin and are associated with synaptic activity. Filamin A modulates ion channel abundance, controls aspects of membrane trafficking, and is involved in the hyperphosphorylation of tau through amyloid signaling (Noam et al., 2014; Burns and Wang, 2017). Differential splicing of filamin A occurs at the same intronic location and to a similar degree in both ZCCHC17 knockdown and AD contexts (**Figure 2E**). Overall, the overlap in splicing changes between ZCCHC17 knockdown human neurons and AD human brains, combined with our previous study demonstrating that ZCCHC17 is reduced in AD brains (Tomljanovic et al., 2018), suggests that loss of ZCCHC17 function may explain a subset of splicing changes seen in AD brain tissue.

### ZCCHC17 Knockdown Alters Nonsense-Mediated Decay

As noted in **Figure 1**, ZCCHC17 also binds to 47 proteins involved in nonsense-mediated decay (NMD). Although NMD has not been extensively examined in the context of AD, we investigated NMD in neurons following ZCCHC17 knockdown. Differential isoform fraction (dIF) was quantified for isoforms with premature termination codons (PTCs) and those without. ZCCHC17 knockdown resulted in significantly increased (p = 1.2 × 10^−11^ by two-sided t-test) expression of PTC-containing isoforms (which are sensitive to NMD), compared to isoforms lacking PTCs (**Figure 3A**). More precisely, there is a sign skew in dIF by PTC status (p = 3.5 × 10^−16^ by Fisher’s exact test). This relationship persists when considering only isoforms which exhibited individually-significant dIF changes upon ZCCHC17 knockdown (p = 2.8 × 10^−3^ by two-sided t-test and p = 1.7 × 10^−4^ by Fisher’s exact test, **Figure 3B**). **Figure 3C** illustrates an isoform switch in *VMP1*, a critical regulatory protein involved in the autophagy process that mediates autophagosome assembly via its ER contacts and maintains neuronal homeostasis (Zhao et al., 2017; Wang et al., 2020; Wang et al., 2021b), which has both NMD-insensitive and NMD-sensitive isoforms. Upon ZCCHC17 knockdown (“siZCC”), overall expression of *VMP1* decreases significantly and expression of the more stable NMD-insensitive isoforms also trends downward. In conjunction, expression of the NMD-sensitive isoform significantly increases, and the isoform fractions converge toward more similar expression levels. Additionally, the C-terminal domain of VMP1, which interacts with beclin-1 and is essential for its autophagy-promoting behavior (Vaccaro et al., 2008), is not present in the NMD-sensitive isoform. NMD-sensitive isoforms are more likely to undergo nonsense-mediated decay and therefore less likely to be translated into functional proteins; thus, a shift toward NMD-sensitive isoforms for a specific gene could result in functional downregulation of that gene.

**Fig. 3.**
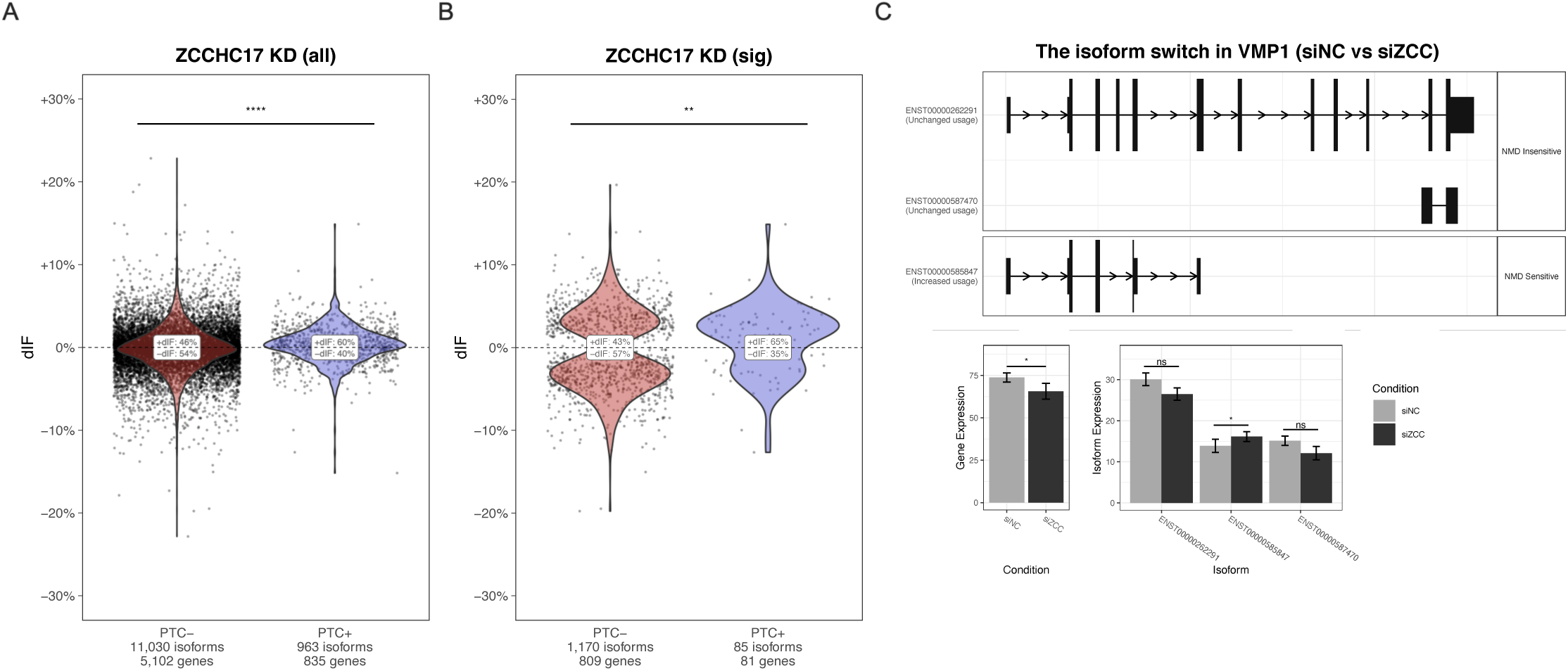
Nonsense-Mediated Decay is Significantly Altered Following ZCCHC17 Knockdown. **(A)** Isoforms containing premature termination codons (PTCs) exhibit significantly increased (p = 1.2 × 10^−11^) differential isoform fraction (dIF) in ZCCHC17 knockdown iPSC-derived neurons differentiated for 42 days post-NPC compared to isoforms lacking PTCs. (**B**) Among the subset of isoforms exhibiting significant changes in fractional expression following ZCCHC17 knockdown, there remains a significant increase (p = 2.8 × 10^−3^) in dIF in PTC+ vs PTC-isoforms. (**C**) VMP1 undergoes an isoform switch in ZCCHC17 knockdown neurons, exhibiting decreased expression of NMD-insensitive isoforms (which lack a PTC) and significantly increased expression (adjusted p = 1.4 × 10^−2^) of its NMD-sensitive isoform (which contains a PTC).

We also investigated NMD in the ROSMAP RNA-seq data, but no significant change was detected in dIF values between NMD-sensitive and NMD-insensitive isoforms (**Supplemental Figure S4B**). Note that below, we also examine NMD in a tau overexpression model, which does show a weak but significant shift in NMD (**Supplemental Figure S4A**). Our overall impression from these data is that, while ZCCHC17 knockdown and tau overexpression do cause a shift in NMD, the present analysis does not support a role for NMD shift in AD brain tissue.

### ZCCHC17 Expression Correlates with Cognitive Resilience

To identify whether ZCCHC17 levels may affect patient outcomes and disease progression in real-world data, we analyzed ZCCHC17 expression data in brain tissue samples from the ROSMAP study and compared its relationship to various cognitive metrics and the accumulation of AD-relevant neuropathologies. After filtering for sufficient RNA quality, 680 donor samples were analyzed from persons exhibiting varying degrees of cognitive decline and disease pathology. ZCCHC17 expression was found to correlate significantly and positively with all listed measures of cognitive function. Notably, these significant relationships were maintained after additionally controlling for various neuropathologies (**Figure 4A**), although with a slight decrease in significance, suggesting that ZCCHC17 mRNA levels in general are minimally affected by increasing AD pathology. Apolipoprotein E4 (APOE4) is a major genetic risk factor of AD, with carriers of even one APOE4 allele having triple the risk of developing late-onset AD compared to patients without it (Uddin et al., 2019). As this patient cohort included a significant number of patients carrying at least one APOE4 allele (APOE4+), we also analyzed this data after splitting samples into groups based on APOE4 status to better understand the impact of this risk factor (**Figure 4B-C**). ZCCHC17 expression was found to correlate significantly with all cognitive metrics in both APOE4+ and APOE4-groups. When pathology was controlled for in the APOE4+ group, this effect was lost for semantic memory, working memory, perceptual orientation, and perceptual speed, and attenuated for episodic memory and global cognition metrics (**Figure 4B**). Controlling for pathology in the APOE4-patient group had less of an effect on the relationship between ZCCHC17 expression and cognition, which was maintained across the majority of cognitive metrics (**Figure 4C**). Global cognition and episodic memory were consistently correlated with ZCCHC17 expression across all sample types and subgroups.

**Fig. 4.**
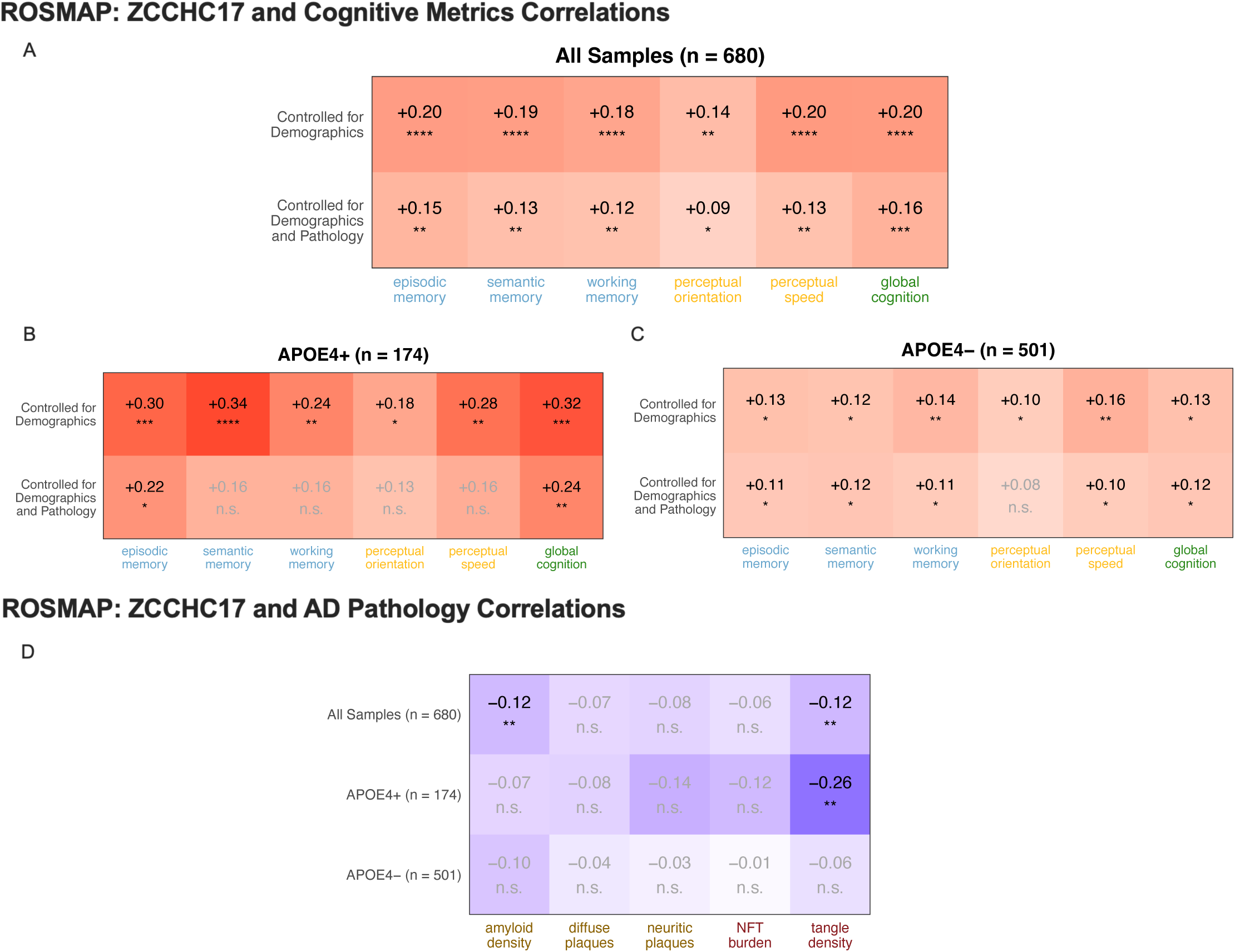
ZCCHC17 Gene Expression Correlates with Cognitive Metrics and AD Pathology. **(A)** ZCCHC17 expression correlates with cognitive metrics in ROSMAP DLPFC samples, whether these metrics are controlled for demographics alone or for both demographics and various measures of neuropathology. (**B-C**) Several cognitive metrics correlate strongly with ZCCHC17 expression in patients carrying an APOE4 allele (APOE4+) and moderately in patients without an APOE4 allele (APOE4-). (**D**) ZCCHC17 expression correlates weakly with amyloid density and tangle density in the full cohort, and strongly with tangle density in APOE4+ subjects. All values shown in heatmaps are Spearman correlation rho values, with red denoting positive correlations and blue denoting negative correlations, and the BH-adjusted significance of each correlation is indicated (* p < 0.05; ** p < 0.01; *** p < 0.001; **** p < 0.0001; n.s., not significant).

Since we observed an effect of pathology on ZCCHC17 correlations with cognition in APOE4+ subjects, we next correlated ZCCHC17 expression with various measures of pathology. As seen in **Figure 4D**, ZCCHC17 expression weakly and negatively correlates with beta-amyloid load in the midfrontal cortex, and the effect is lost when subjects are divided by APOE status. Interestingly, tangle density shows a strong negative correlation with ZCCHC17 expression that is specific for the APOE4+ group. Tau pathology has a well-known effect on cognition (Brier et al., 2016; Hansson et al., 2017), and this explains in part the relative loss of ZCCHC17 correlation with cognitive decline in APOE4+ subjects after regressing out the effect of various neuropathologies. It is unclear why this effect is APOE4-dependent, although several groups have demonstrated that APOE4 exacerbates neurodegeneration and neuronal death in the setting of tau pathology (Shi et al., 2017; Zhao et al., 2020). Whether this effect is mediated in part through ZCCHC17 transcriptional suppression should be investigated in future work (see Discussion).

In summary, the results shown in Figure 4 suggest that cognitive resilience in AD is linked to ZCCHC17 expression levels, and that this effect persists even after controlling for multiple neuropathologies (although APOE4 patients have a more complex picture). We previously found that ZCCHC17 protein levels decline early in the course of AD, before significant astrogliosis or neuronal loss (Tomljanovic et al., 2018). Taken together, one possible interpretation of these findings is that ZCCHC17 supports cognitive function in part through maintenance of RNA splicing/processing, and declines early in AD at the protein level for reasons other than transcriptional suppression. This loss then contributes to synaptic dysfunction and cognitive impairment by impeding normal RNA splicing and processing mechanisms.

The implication of these data is that elevated baseline ZCCHC17 expression levels may support cognitive resilience in the setting of progressive AD pathology. To test this hypothesis further, we used PrediXcan (Gamazon et al., 2015) to quantify predicted levels of ZCCHC17 expression, leveraging the GTEx database to build a model which we then applied to GWAS data. This methodology enables the use of GWAS/gene expression relationships across diverse tissues in a training set to predict genetically-regulated levels of gene expression in a given tissue from subsequent GWAS data (Hohman et al., 2017; Dumitrescu et al., 2020). Higher predicted expression of ZCCHC17 across 15 tissues was associated with higher cognitive resilience (GBJ p = 0.02), including associations in heart tissue (atrial appendage β = 0.17, p = 0.005) and brain tissue (putamen β = 0.09, p = 0.002) that remained significant after correction for multiple comparisons. Results were slightly stronger when participants with AD were removed from the analytical models (GBJ p = 0.004), revealing additional associations in tibial nerve (β = 0.22, p = 0.001, **Figure 5A**), thyroid (β = 0.15, p = 0.003), pituitary (β = 0.09, p = 0.01), and colon (β = 0.10, p = 0.02) after correction for multiple comparisons. When assessing the validity of these significant predicted expression models in ROSMAP, higher predicted expression in 5/6 tissues that related to resilience also showed associations with higher measured ZCCHC17 expression in the brain (**Figure 5B-E**), suggesting that even the predicted expression models built from peripheral tissues have relevance to the brain. Notably, predicted expression in heart, brain, nerve, colon, and thyroid was significantly associated with both observed expression in ROSMAP and resilience. Taken together, the above analysis further supports the hypothesis that elevated ZCCHC17 levels are neuroprotective in the setting of AD pathology.

**Fig. 5.**
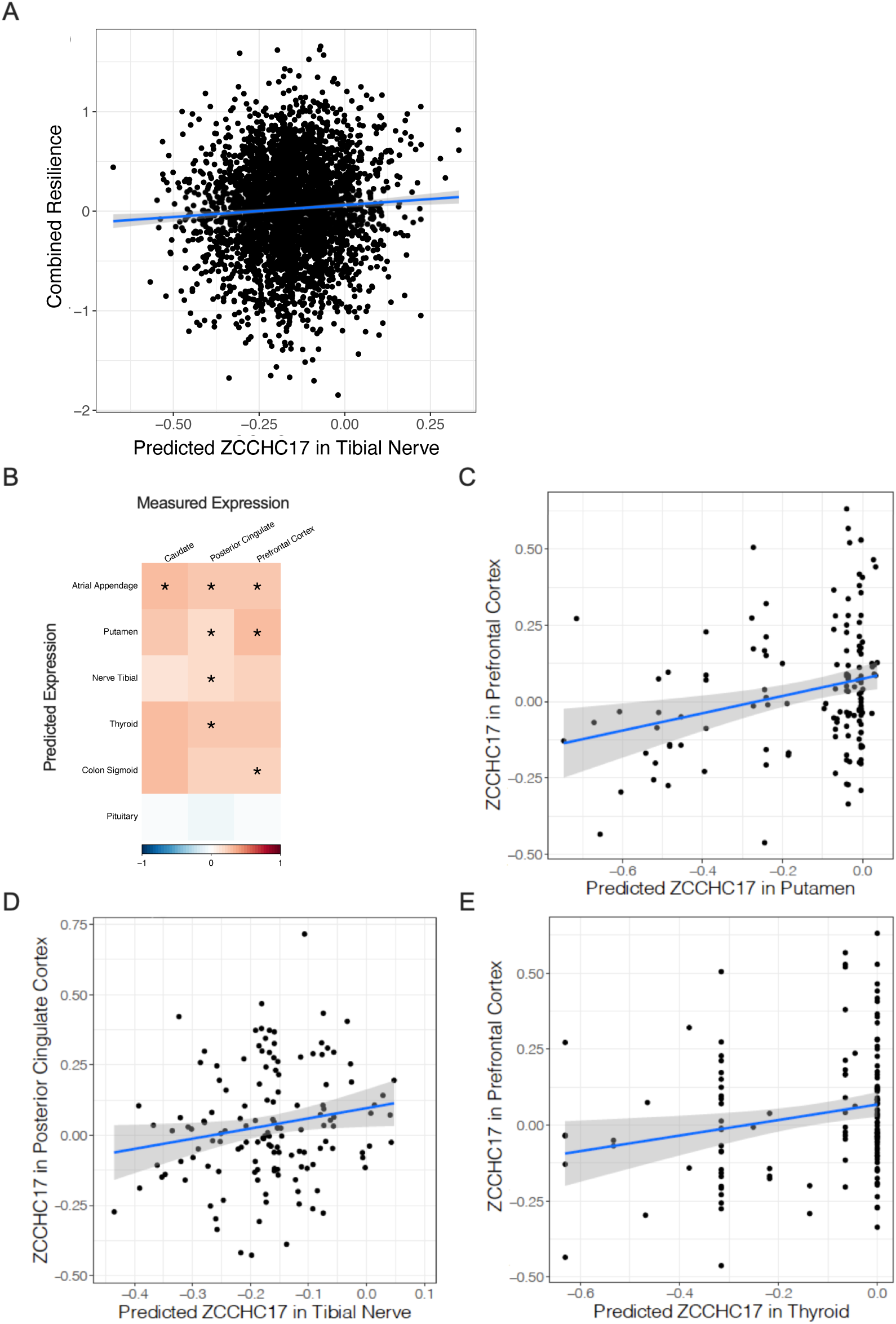
Predicted Expression of ZCCHC17 Correlates with Resilience in Large GWAS Study and Observed Expression in Brain Tissue. **(A)** Higher ZCCHC17 predicted expression is associated with better-than-predicted cognitive performance given an individual’s amyloid burden, or resilience. Regression line shading indicates standard error of measurement. To validate our genetically-predicted expression models, panels B-E demonstrate association between predicted expression built in GTEx and observed expression measured in ROSMAP. (**B**) Correlation plot for all tissues that showed a significant association between predicted expression of ZCCHC17 and resilience. Color bar represents Pearson’s correlation coefficient (* p < 0.05). (**C-E**) Predicted expression of ZCCHC17 is presented along the x-axis, and observed expression is presented along the y-axis.

Although ZCCHC17 has not been identified as an AD-associated risk gene through prior genome-wide SNP analysis, the above findings motivated us to determine whether there are any AD-risk variants in the ZCCHC17 loci that are nominally associated with AD, and if so, the level of significance of this association. We analyzed the summary statistics from a previously published AD GWAS (71,880 AD cases and 383,378 controls) (Jansen et al., 2019) to identify SNPs in ZCCHC17 that are associated with clinical Alzheimer’s Disease. Two variants (rs59705505 at 3.73 × 10^−6^ and rs11336043 at 9.24 × 10^−6^) are significant loci at *P* < 1 × 10^−5^. While not genome-wide significant at the widely used threshold of 5 × 10^−8^, these variants are nominally significant in the NHGRI-EBI GWAS Catalog (Buniello et al., 2019). Although intriguing, it should be noted that ZCCHC17 is in a gene-dense region of the genome, and since these SNPs are not located within ZCCHC17, it is difficult to assign them unambiguously to ZCCHC17.

### ZCCHC17 and Tau Share Common Binding Partners

Tau has been shown to bind to DNA and play a role in transcriptional regulation (Greenwood and Johnson, 1995; Hua et al., 2003). Tau also binds to RNA (Schröder et al., 1984; Dinkel et al., 2015) and tau overexpression has been linked to splicing abnormalities (Apicco et al., 2019; Hsieh et al., 2019). To investigate whether tau overexpression may play a causal role in ZCCHC17 dysfunction, ZCCHC17 binding partners were compared to published tau interactors in human postmortem brain tissue (Hsieh et al., 2019) (**Supplemental Figure S5**). ZCCHC17 binding partners are significantly enriched (p = 7.0 × 10^−14^ by Fisher’s exact test using the brain proteome as background (Johnson et al., 2020)) for tau binding partners, with 55% (50 proteins) of ZCCHC17 interactors being shared with tau (**Supplemental Figure S5A**). These shared interactors include all synaptic vesicle proteins that interact with ZCCHC17, 42% of ZCCHC17 RNA splicing proteins, 57% of ZCCHC17 RNA binding proteins, and 66% of ZCCHC17 nonsense-mediated decay proteins. When looking specifically at tau interactors in AD brains, ZCCHC17 binding partners are also significantly enriched (p = 3.7 × 10^−10^ by Fisher’s exact test) for tau interactors, with 33% (30 proteins) of ZCCHC17 interactors shared with tau under AD conditions (**Supplemental Figure S5B**). These shared interactors include all synaptic vesicle proteins that interact with ZCCHC17, 25% of ZCCHC17 RNA splicing proteins, 32% of ZCCHC17 RNA binding proteins, and 38% of ZCCHC17 nonsense-mediated decay proteins.

### Tau Overexpression and ZCCHC17 Knockdown in iPSC-Derived Neurons Induce Similar RNA Processing Abnormalities

ZCCHC17 protein binding partners intersect with those of tau, suggesting that they may share overlapping functional roles in neurons. To investigate the effects of alterations in expression of tau on the generation of functional proteins and RNA, differential splicing analysis was conducted on previously-published gene expression data from a tau overexpression model in human iPSC-derived neurons (Raj et al., 2018). 99 differentially spliced clusters, corresponding to 91 unique DSGs, were identified when comparing tau overexpression samples with empty vector control samples (**Figure 6A, Supplemental Table S3**). Of the 91 DSGs induced by tau overexpression, 17 were also affected following ZCCHC17 knockdown (**Figure 6B**). GO analysis of these 17 shared genes revealed significant enrichment in only 2 pathways: “cytoskeleton” (strength = 4.63, adjusted p-value = 6.9 × 10^−2^) and “actin cytoskeleton” (strength = 11.76, adjusted p-value = 4.8 × 10^−2^). 13 out of the 91 tau overexpression DSGs were also affected in AD postmortem tissue from ROSMAP (**Supplemental Table S3**). Seven genes were significantly differentially spliced in all three data sets examined: ACTN4, AGRN, MYO6, NEO1, SEPTIN9, TACC2, and TSC22D3 (**Supplemental Table S3**). α-Actinin-4 (ACTN4) is a Ca^2+^-sensitive actin-binding protein that promotes the remodeling of dendritic spines by metabotropic glutamate receptors (Walikonis et al., 2001; Kalinowska et al., 2015). The significant effect on splicing of ACTN4 in ZCCHC17 knockdown neurons stems from significant changes in clusters 3931 (adjusted p-value = 1.2 × 10^−2^) and 3932 (adjusted p-vale = 2.5 × 10^−2^), with a greater effect size (loglr = 5.9) in cluster 3931 (**Figure 6C**). Tau overexpression neurons exhibited a significant splicing change in ACTN4 at the intron location matching cluster 3931 (adjusted p-value = 4.2 × 10^−2^), with a similar effect size (loglr = 5.4) (**Figure 6D**). ACTN4 is also a significant DSG in the ROSMAP-AD analysis, with high effect size for its affected clusters (loglr = 7.1 and 5.3), but these clusters correspond to unique intron locations in the ROSMAP data set and are not directly comparable.

**Fig. 6.**
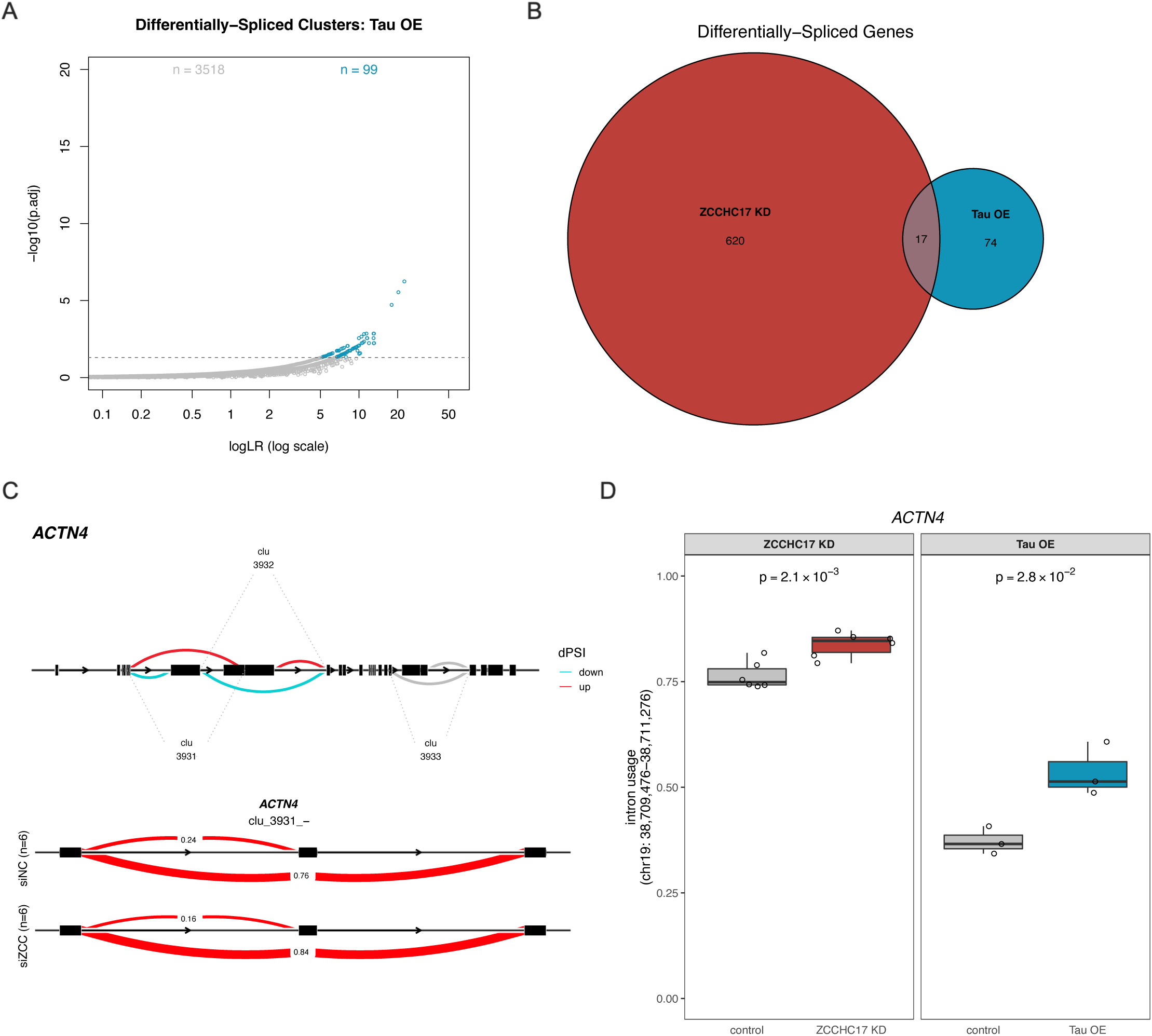
Tau Overexpression Induces RNA Splicing and Processing Alterations Similar to ZCCHC17 Knockdown-Dependent Changes. **(A)** 99 intron clusters exhibited differential splicing between tau overexpression iPSC-derived neurons (n = 3) and controls (n = 3). (**B**) 17 DSGs are shared between ZCCHC17 knockdown neurons (“ZCCHC17 KD,” 637 DSGs) and tau overexpression neurons (“Tau OE,” 91 DSGs). (**C**) In ZCCHC17 knockdown neurons, ACTN4 is differentially spliced at clusters 3931 (adjusted p = 1.2 × 10^−2^) and 3932 (adjusted p = 2.5 × 10^−2^), with significantly decreasing dPSI in blue and significantly increasing dPSI in red. Other clusters identified by LeafCutter for ACTN4 are shown in grey. The alteration in cluster 3931 corresponds to an exon skipping event, with usage proportions of significantly-changing introns shown in red in the lower panel. Exons are shown in black. (**D**) Alternative splicing occurs at the same location and to a similar degree in both ZCCHC17 knockdown neurons and tau overexpression neurons, with significant changes in both (p = 2.1 × 10^−3^ and 2.8 × 10^−2^, respectively).

Since tau binding partners included a large number of NMD proteins (**Supplemental Figure S5**), we also investigated the NMD mechanism in the tau overexpression data. Although the effect was weaker than observed in ZCCHC17 knockdown neurons, tau overexpression neurons exhibited a significant increase in dIF for PTC-containing isoforms (p = 6.6 × 10^−3^ by two-sided t-test) as well as a significant positive skew in dIF for PTC-containing isoforms (p = 0.03 by Fisher’s exact test), indicating a similar shift in RNA processing (**Supplemental Figure S4A**). This previously unreported effect of tau overexpression on RNA processing, while weak, suggests that tau may be affecting RNA metabolism through multiple mechanisms related to RNA splicing. As noted above, we also investigated this mechanism in the ROSMAP AD data set, but no significant change was detected in dIF values between NMD-sensitive and NMD-insensitive isoforms (**Supplemental Figure S4B**), suggesting that if NMD changes are relevant in AD, their effects are either weak or difficult to detect in autopsy brain tissue.

## Discussion

In this study, we have further explored the function of ZCCHC17, originally identified by our group as a transcriptional regulator whose dysfunction in AD may contribute to dysregulation of synaptic gene expression (Tomljanovic et al., 2018). We define ZCCHC17’s interactions at the RNA processing and protein level and compare its impacts across multiple disease-relevant models. Our co-immunoprecipitation study provides a comprehensive list of proteins which interact with ZCCHC17 in human neurons under basal conditions, and defines relevant groups of interactions based on protein function. The network of proteins which bind to ZCCHC17 includes synaptic proteins as well as many involved in the regulation and stability of RNA and downstream proteins, specifically from RNA splicing, RNA binding, and nonsense-mediated decay pathways. RNA metabolism proteins have been implicated in several neurodegenerative diseases; amyotrophic lateral sclerosis (ALS) and autism have been linked to the dysfunction of RNA-binding proteins (Parikshak et al., 2016; Kapeli et al., 2017), and pathologic RNA-protein aggregates have been observed in ALS and inclusion body myopathy (Ramaswami et al., 2013; Taylor et al., 2016). In late-onset AD, tau has been shown to interact with RNA (Schröder et al., 1984; Dinkel et al., 2015), RNA-binding proteins (Broccolini et al., 2000; Bai et al., 2013; Gunawardana et al., 2015; Hsieh et al., 2019), and the ribosome (Ding, 2005; Koren et al., 2020). ZCCHC17 has previously been found to interact with other transcriptional regulators (Chang et al., 2003) as well as splicing factors (Ouyang, 2009; Lin et al., 2017), but our work directly shows these interactions in human neurons and provides a more comprehensive view of its interactions with other proteins.

We evaluated and quantified differential splicing in human neurons following ZCCHC17 knockdown and compared these alterations to those observed in brain tissue from AD patients. We show that ZCCHC17 knockdown leads to dysregulation of mRNA splicing across a broad range of categories (including synaptic genes) and that these splicing abnormalities significantly overlap with splicing abnormalities in AD brain tissue. Pre-mRNA splicing plays a major role in the development of many human diseases; at least 20-30% of disease-causing mutations are associated with aberrant splicing (Wang and Cooper, 2007; Lim et al., 2011). Neurodegenerative diseases in particular have been linked to specific splicing alterations, including *MAPT* exon 10 splice site mutations in frontotemporal dementia and parkinsonism linked to chromosome 17 (FTDP-17) (Hutton et al., 1998; Trabzuni et al., 2012) and potentially *SCNA* isoforms in Parkinson’s Disease (Oueslati, 2016; Kaji et al., 2020). In late-onset AD, intron retention increases to levels significantly beyond those observed in physiological aging, and widespread disruption of splicing mechanisms alters the brain transcriptome (Faustino, 2003; Vaquero-Garcia et al., 2016; Raj et al., 2018; Adusumalli et al., 2019). The DSGs in our ZCCHC17 knockdown model which overlap with those in AD brain tissues are enriched for synaptic genes, further supporting the view that loss of ZCCHC17 function in AD may contribute to synaptic dysfunction.

Alternative splicing is a complex phenomenon that alters RNA and protein magnitude and diversity. The downstream effects of dysregulated alternative splicing are likely much broader and of a greater magnitude when splicing defects significantly disrupt RNA and protein expression for a prolonged period of time; for example, over the decades of pathologic progression which precede clinical onset of AD (Perrin et al., 2009; Jack, 2011). How ZCCHC17 dysfunction interacts with ongoing RNA metabolic defects in AD and how this contributes to cognitive dysfunction is an interesting question. Indeed, ZCCHC17 is a relatively unstudied protein, and the work detailed here elevates the importance of further investigating its role in health and disease. As noted in one of the original papers that discovered ZCCHC17, the protein contains a number of potential phosphorylation sites, including 10 for protein kinase C, 6 for casein kinase II, and 2 for cAMP-dependent kinase (Chang et al., 2003). Although early work found that cytoplasmic ZCCHC17 co-fractionates with ribosomes (Gueydan et al., 2002), recent studies have focused on its nuclear function and revealed that it possesses a diverse range of functions relevant to both mRNA (Lin et al., 2017) and rRNA (Lin et al., 2019) processing. This has led some to suggest that ZCCHC17 may play a broad role in coordinating and maintaining cell homeostasis (Lin et al., 2017).

The effect of ZCCHC17 expression on cognition is both interesting and understandable given these wide-ranging effects on neuronal health. Our group previously showed that ZCCHC17 protein levels decline early in the course of AD, before significant gliosis or neuronal loss (Tomljanovic et al., 2018). Interestingly, we show here that ZCCHC17 mRNA levels, while sensitive to some aspects of AD pathology, are for the most part resilient to AD pathology in patients who lack the APOE4 allele, and correlate with a broad range of cognitive metrics even after controlling for neuropathology. One possible interpretation of these data is that the level of functional ZCCHC17 protein declines in AD, and higher ZCCHC17 expression can partially buffer this effect. This interpretation is further supported by our PrediXcan analysis of ZCCHC17 expression, which shows that predicted levels of ZCCHC17 correlate with cognitive resilience in patients with documented AD pathology on imaging. Although we have presented evidence supporting an impact of ZCCHC17 on cognitive function in this study, the exact mechanism for this impact is not clear. While ZCCHC17’s role in regulating synaptic gene expression is an obvious candidate, the broader influence that ZCCHC17 may exert on neuronal health should not be ignored, and ZCCHC17 loss of function may contribute to neurodegeneration through a variety of mechanisms. Indeed, while in this paper we have focused on ZCCHC17 interactors involved in RNA processing (which comprise the majority of the interactors we identified), we also found that proteins involved in synaptic vesicle cycling, autophagy, and stress granule formation interact with ZCCHC17. We have not explored the relevance of these findings in this study, but they warrant further investigation in future studies of ZCCHC17 and its contribution to neurodegeneration.

We also uncovered a potential interaction between ZCCHC17 and tau, initially by investigating the overlap of protein binding partners via our co-IP study, which suggests that tau accumulation in AD could disrupt functionally relevant protein binding partners of ZCCHC17. A hypothesized sequestration of the network of proteins supporting ZCCHC17 function by AD-related tau would likely interfere with its ability to regulate RNA processing, as >70% of these overlapping proteins are involved in RNA splicing, RNA binding, or nonsense-mediated decay pathways. Our follow-up alternative splicing and NMD analysis found similarities between our ZCCHC17 knockdown model and published tau overexpression data at the RNA processing level, which may reflect downstream effects of this protein-level relationship. Future work will need to examine whether ZCCHC17 function is affected by tau in AD brain tissue through sequestration of ZCCHC17 interactors. Interestingly, we also found a strong correlation between ZCCHC17 gene expression and neurofibrillary tangle burden in patients carrying an APOE4 allele, which also impacts the link between ZCCHC17 expression and cognition in APOE4 carriers (particularly semantic memory, working memory, and perceptual memory) after regressing out the effect of AD pathology. Additional investigation is necessary to further elucidate the relationship between ZCCHC17, tau, and APOE.

## Abbreviations

AD: Alzheimer’s disease
NPC: neural progenitor cells
PBS: phosphate buffered saline
SDS-PAGE: sodium dodecyl sulfate–polyacrylamide gel electrophoresis
TBST: Tris-buffered saline with 0.1% Tween
MS: mass spectrometry
FDR: false discovery rate
hiPSC: human induced pluripotent stem cell
ROSMAP: Rush Memory and Aging Project
DSGs: Differentially spliced genes
GO: Gene ontology
PTCs: Premature termination codons
RIN: RNA integrity number
PMI: post mortem interval
NMD: nonsense mediated decay
dIF: differential isoform fraction
BH: Benjamini-Hochberg
GWAS: genome-wide association study
DEGs: differently expressed genes
DLPFC: dorsolateral prefrontal cortex
GBJ: generalized Berk–Jones

## Declarations

### Ethics approval and Consent to participate

All human cell line work was performed on de-identified cell lines and approved by the Columbia University Institutional Review Board (IRB).

### Availability of supporting data

Newly-generated RNA-seq data have been deposited in GEO (https://www.ncbi.nlm.nih.gov/geo/, GSE199241) and are publicly available as of the date of publication. ROSMAP RNA-seq data is available via the AMP-AD data portal through Synapse (https://www.synapse.org/#!Synapse:syn3219045) and the RADC Research Resource Sharing Hub (https://www.radc.rush.edu).

## Acknowledgements

Not Applicable

---

**Fig. S1.**
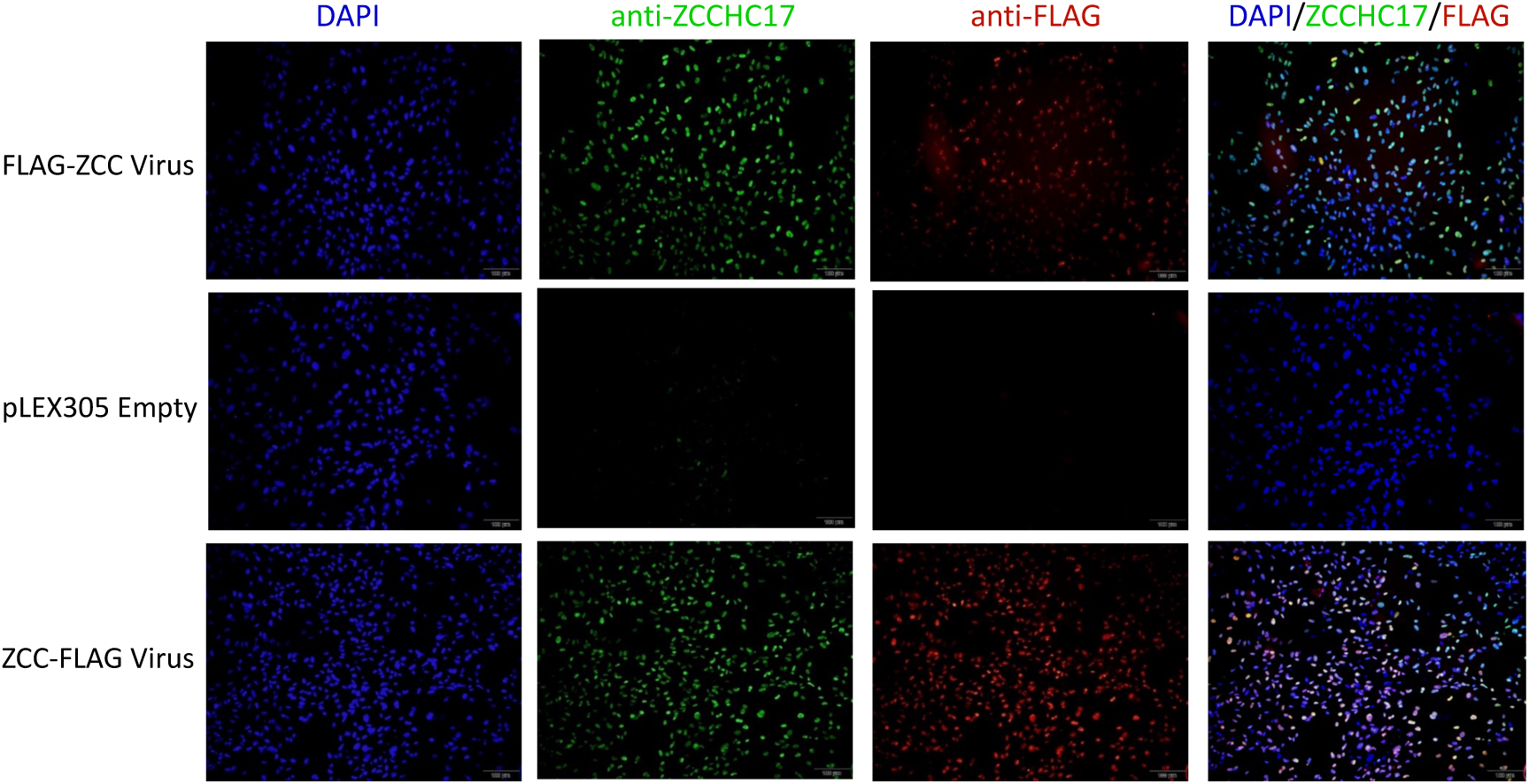
Neurons Infected with FLAG-tagged ZCCHC17 Viral Constructs Express Both FLAG and Increased Levels of ZCCHC17. Immunocytochemistry images from 52-day differentiation of iPSC-derived neurons stained with antibodies against ZCCHC17 and FLAG. Control neurons infected with pLEX_305 empty viral vector do not express FLAG and express ZCCHC17 at baseline control levels.

**Fig. S2.**
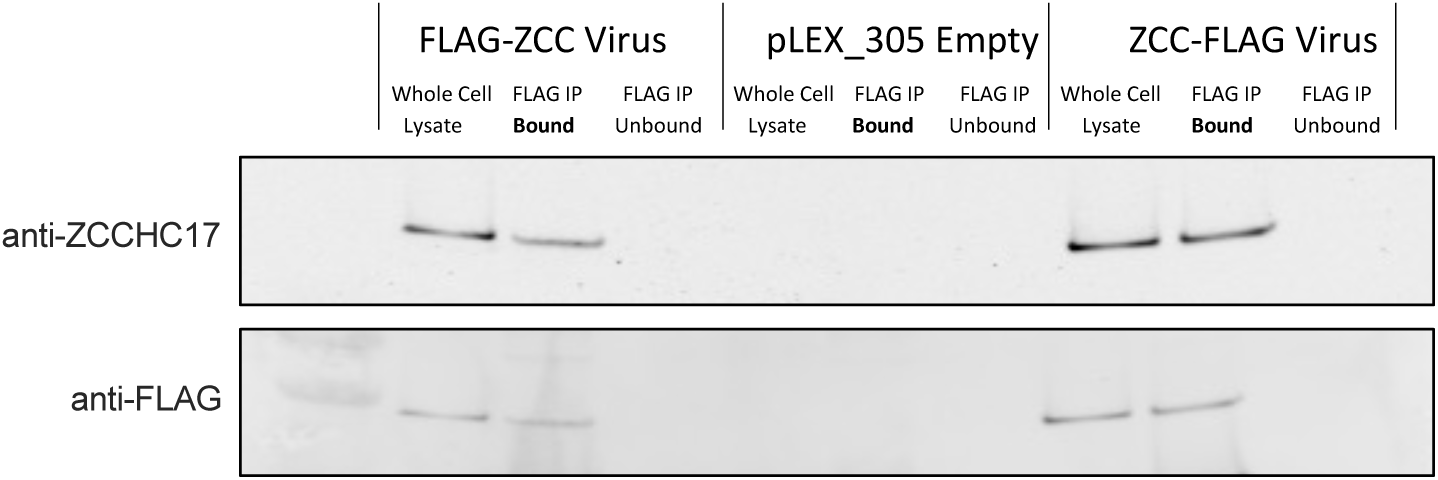
FLAG-tagged ZCCHC17 was Immunoprecipitated from Neurons Prior to LC-MS. Immunoblot of whole cell lysate, fraction separated by anti-FLAG immunoprecipitation (“FLAG IP Bound”), and remaining depleted lysate (“FLAG IP Unbound”) from 68-day differentiation of human iPSC-derived neurons infected with FLAG-tagged ZCCHC17 viral construct or pLEX_305 empty viral vector. ZCCHC17 and FLAG were detected in the anti-FLAG bound fraction after immunoprecipitation from FLAG-expressing neurons and not from control neurons.

**Fig. S3.**
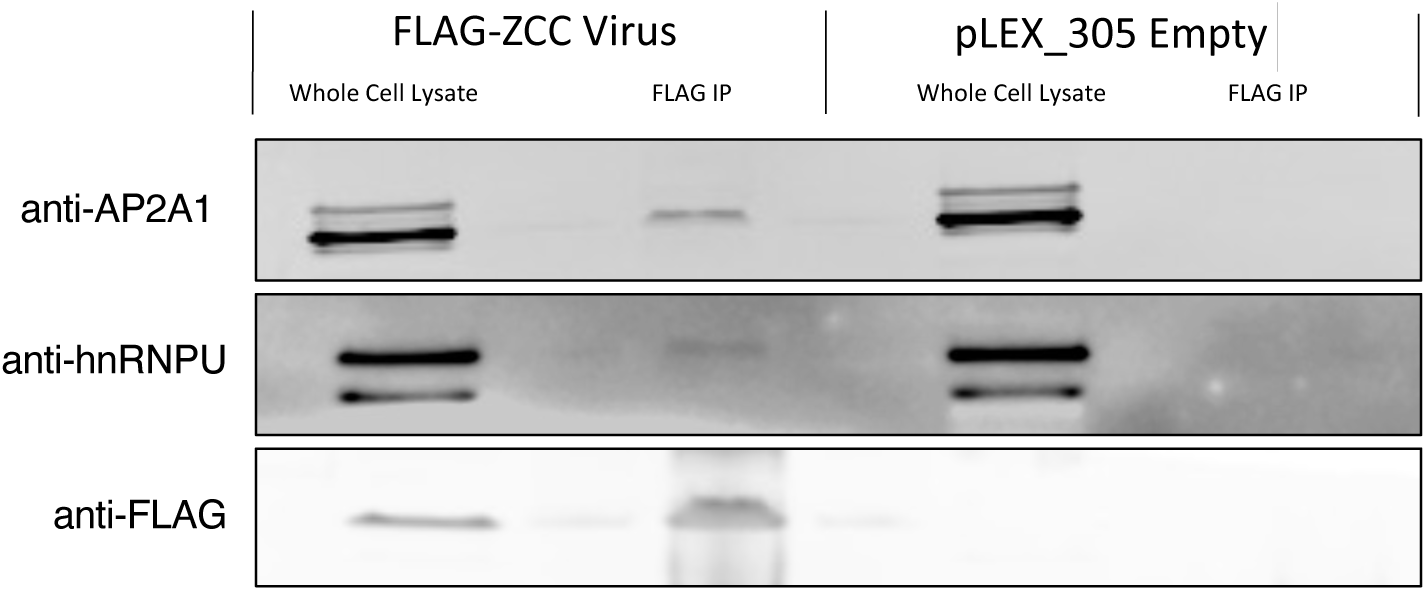
Binding Partners of ZCCHC17 Confirmed by Western Blot. Immunoblot of whole cell lysate and fraction separated by anti-FLAG immunoprecipitation (“FLAG IP”) from 70-day differentiation of human iPSC-derived neurons infected with FLAG-tagged ZCCHC17 viral constructs or pLEX_305 empty viral vector. AP2A1 and hnRNPU are present in the anti-FLAG bound fraction after immunoprecipitation from FLAG-expressing neurons and not from control neurons.

**Fig. S4.**
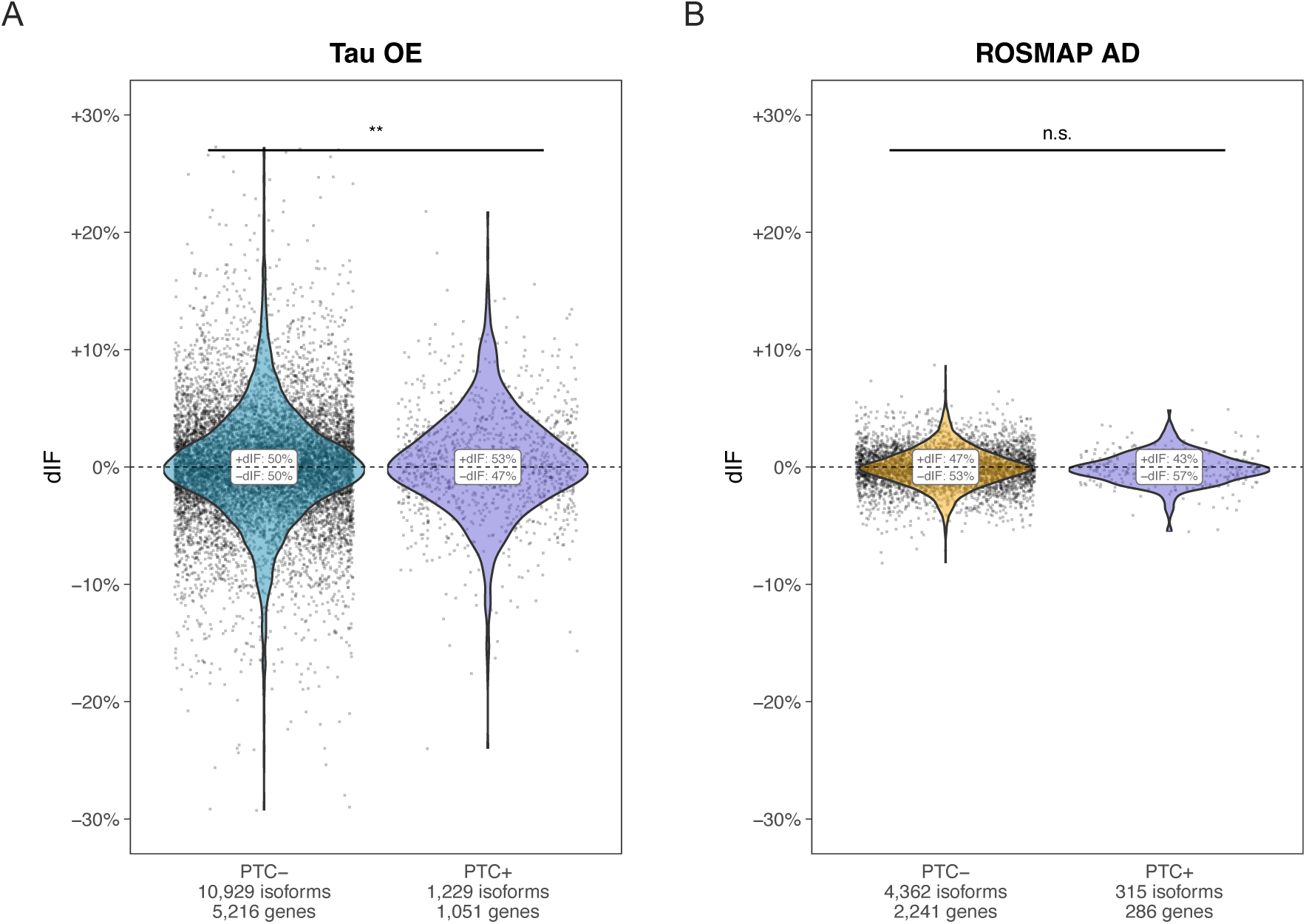
NMD-Sensitive Differential Isoform Fraction Changes Significantly in Tau Overexpressing Neurons but not in ROSMAP AD Data. **(A)** Nonsense-mediated decay is altered following tau overexpression, with a significant increase (p = 6.6 × 10^−3^) in dIF among isoforms with annotated premature termination codons (PTCs) relative to isoforms lacking PTCs. (**B**) In AD human tissue samples from ROSMAP, differential isoform fraction (dIF) is not significantly increased for isoforms containing PTCs relative to isoforms lacking PTCs.

**Fig. S5.**
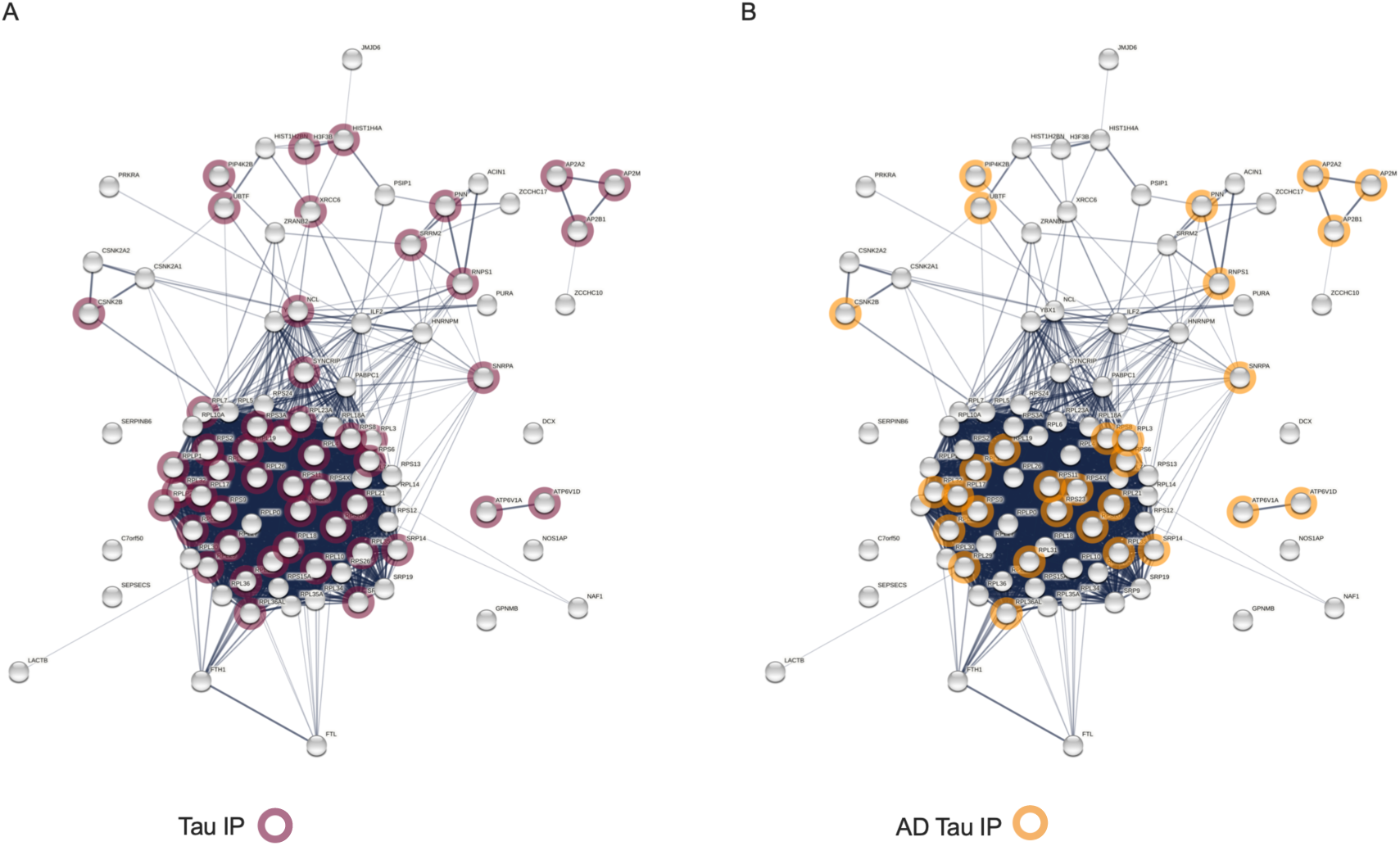
ZCCHC17 and Tau Share a Significant Number of Binding Partners. **(A)** Binding partners of ZCCHC17 are significantly enriched for tau binding partners. Previously-published tau immunoprecipitation data from human postmortem brain tissue **[1]** is shown, with 50/91 (55%, p = 7.0 × 10^−14^ by Fisher’s exact test) ZCCHC17 binding partners identified as tau binding partners. (**B**) 30/91 (33%, p = 3.7 × 10^−10^ by Fisher’s exact test) ZCCHC17 binding partners are identified as AD-associated tau binding partners.

**Table S1 (separate file) List of Proteins Immunoprecipitated with ZCCHC17**

**Table S2 (separate file) List of Differentially Expressed Genes in ZCCHC17 Knockdown Neurons**

**Table S3 (separate file) List of Differentially Spliced Genes in ZCCHC17 Knockdown Neurons, ROSMAP-AD Brain Tissue, Tau-OE Neurons, and Enriched Gene Ontology Groups**

